# Influence of prior beliefs on perception in early psychosis: effects of illness stage and hierarchical level of belief

**DOI:** 10.1101/421891

**Authors:** J. Haarsma, F. Knolle, J.D. Griffin, H. Taverne, M. Mada, I.M. Goodyer, the NSPN Consortium, P.C. Fletcher, G.K. Murray

## Abstract

Alterations in the balance between prior expectations and sensory evidence may account for faulty perceptions and inferences leading to psychosis. However, uncertainties remain about the nature of altered prior expectations and the degree to which they vary with the emergence of psychosis. We explored how expectations arising at two different levels – cognitive and perceptual – influenced processing of sensory information and whether relative influences of higher and lower level priors differed across people with prodromal symptoms and those with psychotic illness. In two complementary auditory perception experiments, 91 participants (30 with first episode psychosis, 29 at clinical risk for psychosis, and 32 controls) were required to decipher a phoneme within ambiguous auditory input. Expectations were generated in two ways: an accompanying visual input of lip movements observed during auditory presentation, or through written presentation of a phoneme provided prior to auditory presentation. We determined how these different types of information shaped auditory perceptual experience, how this was altered across the prodromal and established phases of psychosis, and how this relates to cingulate glutamate levels assessed by magnetic resonance spectroscopy. The psychosis group relied more on high level cognitive priors compared to both healthy controls and those at clinical risk for psychosis, and more on low level perceptual priors than the clinical risk group. The risk group were marginally less reliant on low level perceptual priors than controls. The results are consistent with previous theory that influences of prior expectations in psychosis in perception differ according to level of prior and illness phase.

**General scientific summary:** What we perceive and believe on any given moment will allow us to form expectations about what we will experience in the next. In psychosis, it is believed that the influence of these so-called perceptual and cognitive ‘prior’ expectations on perception is altered, thereby giving rise to the symptoms seen in psychosis. However, research thus far has found mixed evidence, some suggesting an increase in the influence of priors and some finding a decrease. Here we test the hypothesis that perceptual and cognitive priors are differentially affected in individuals at-risk for psychosis and individuals with a first episode of psychosis, thereby partially explaining the mixed findings in the literature. We indeed found evidence in favour of this hypothesis, finding weaker perceptual priors in individuals at-risk, but stronger cognitive priors in individuals with first episode psychosis.

## 1.1. Background

It has been hypothesized that the brain forms a model of the world by actively trying to predict it and to update these predictions iteratively by function of the prediction error, a hierarchical computational framework usually referred to as predictive coding (Rao & Ballard et al., 1999; Bar, 2009; Friston, 2005 & 2009; Bastos et al., 2012; Clark et al., 2013 & 2015; Hohwy et al., 2013; Knill et al., 2004). In this framework, the formation of delusional beliefs and hallucinatory experiences are proposed to be due to alterations in the cognitive and biological mechanisms of predictive coding (Fletcher & Frith, 2009; Adams et al., 2013).

Whilst initial clinical studies documenting alterations in the way the expectation influences perception in psychosis are promising in demonstrating case-control alterations in various behavioural measures of predictive coding (eg Shergill et al 2005, Teufel et al., 2010; Powers et al 2017), it is already clear that there will be no straightforward unifying explanation of psychosis in simple terms of priors being “too strong” or “too weak” in general. Predictive processing theory envisions a highly interlinked (cortical) cognitive hierarchy, where different layers aim to predict the incoming input from lower-layers (Rao & Ballard et al., 1999; Bar et al., 2009; Friston, 2005 & 2009; Bastos et al., 2012; Clark et al., 2013 & 2015; Hohwy et al., 2013; Knill et al., 2004). Moving up the hierarchy, the predictions become more abstract, ranging from lower-level sensory prediction to higher-order beliefs about the environment. It therefore does not suffice to ask the question whether prior expectations are stronger or weaker in psychosis. Instead in order to form a complete picture of the underlying mechanisms of psychosis, we need to look at the contribution of different types of prior expectations, including both sensory expectations and higher-level beliefs about the environment.

Recent influential predictive coding accounts of psychosis have emphasized that priors at low and high hierarchical levels may be differentially affected in psychotic illness. For example, Sterzer et al (2018) conclude that “In contrast to weak low-level priors, the effects of more abstract high-level priors may be abnormally strong” in psychosis. This postulate is mainly drawn through a combination of theoretical arguments and synthesis across diverse studies. To our knowledge no single study has yet demonstrated a combination of weak low-level perceptual priors and strong high-level cognitive priors in patients with psychosis, although Schmack (2013) and colleagues provided supportive evidence in a study of individual differences in healthy individuals. Those authors delineated priors at different hierarchical levels by manipulating what they referred to as perceptual priors and cognitive priors in two related experiments; they found that delusional ideation in health (sometimes termed delusion proneness) was associated with a decrease in the contribution of perceptual priors, and an increase in the contribution of cognitive priors, highlighting the importance to separate the two (Schmack et al., 2013). Clearly, clinical studies are required testing the hypothesis of simultaneous weak low-level and strong high-level priors in psychotic illness, yet few have been attempted. One exception was another study from Schmack and colleagues, who found evidence against differential strengths of sensory and cognitive priors in schizophrenia (Schmack et al 2017).

A further complexity is that cognitive and biological mechanisms of psychosis may be markedly different at different illness stages, adding nuance to the attractive, yet arguably overly simplistic, continuum model of psychosis. Previous reviews acknowledge that there may be evolving patterns of cognitive and/or physiological disturbances over time as psychotic illness develops (Fletcher & Frith, 2009; Adams et al., 2013; Heinz et al., 2018). In many cases psychotic illness is heralded by the development of delusions (often delusional interpretations of hallucinations) after a prodromal period of hallucinatory experiences without delusional interpretation and/or delusional mood. In the context of weak low level (sensory) priors and high precision of sensory prediction errors, delusions may emerge as result of compensatory increases in the precision of high-level beliefs (i.e. enhanced high level, cognitive priors) (Adams et al 2013, Sterzer et al 2018, Heinz et al 2018). It follows then that in the very early phases of psychosis, prior to the development of delusions, such compensatory increases in the precision of high level beliefs may be yet to emerge. Although one previous study found alterations in the utilisation of priors in individuals at clinical risk for psychosis (putatively in the prodrome) compared to controls (Teufel et al 2015), this study did not include any patients with established psychotic illness, and thus none of the sample had developed delusions at the time of the experiment. It thus remains unclear whether, or how, alterations in the use of higher or lower level priors changes as psychotic illness emerges.

We acknowledge the vital importance of the range of previous studies exploring the contribution of prior expectation in perception in psychosis. However, here we argue that two important aspects of the predictive coding account have been largely neglected in empirical clinical studies : the contribution of different disease stages to the effect of prior expectations, and the type of prior expectation. It is the aim of the present study to bring these two together, by studying how different prior expectations are affected throughout individuals at risk for psychosis and individuals who recently had an episode of psychosis.

In order to test the hypothesis that sensory and cognitive priors are differently used depending on the stage of psychosis, we designed two novel auditory perception paradigms, one testing the influence of lip-movements on auditory perception (perceptual priors) and a second testing the influence of learned written-word-sound associations on auditory perception (cognitive priors); and we gathered data on these two paradigms in two patient groups - individuals at elevated clinical risk for psychosis, and individuals who recently had their first episode of psychosis, and compared them to a group of healthy controls. Help-seeking individuals who are at-risk for psychosis usually have sub-clinical psychotic symptoms that are not severe or frequent enough to warrant a clinical diagnosis, but are at considerably raised risk of developing a psychotic illness in the short to medium term (Yung et al., 2003). Studying these early stages of illness may help us to understand the mechanisms underlying the emergence of a psychosis by examining which aberrancies precede psychosis and might therefore be predictive of developing psychosis.

The first paradigm (from now on ‘perceptual priors task’) assesses the influence of lip-movements on auditory perception. Lip-movements have been shown to influence auditory perception. McGurk and MacDonald (1976) showed that when individuals where presented with an auditory /Ba phoneme in combination with lip-movements pronouncing /Ga, most individuals perceive a mixture between the two, i.e. /Da. This effect has become known as the McGurk illusion (McGurk & MacDonald, 1976). Studies of the neural mechanisms underlying the influence of lip-movements on auditory perception provide support for the Bayesian framework, in that lip-movements are suggested to constitute a prior expectation with respect to the incoming auditory signal (Arnal et al., 2012; Blank & Davis, 2016). One previous study of mainly male, middle-aged adults with chronic schizophrenia documented a diminishment in perceiving the McGurk illusion, relying more on the auditory input; the finding that was associated with illness chronicity (White et al., 2014). Pearl et al (2009) also studied the McGurk illusion in schizophrenia, finding mixed results: adolescents with schizophrenia, but not adults with schizophrenia, showed a diminished illusory effect. Schizophrenia has been associated with a diminished ability in using lip-movements in aiding auditory discrimination, suggesting aberrancy in the ability to integrate the two sources of information (Myslobodsky et al., 1992; de Gelder et al., 2002; Ross et al., 2007; Szycik et al., 2013). However, it remains unclear whether the influence of prior information in auditory perception is altered in the early stages of psychosis, as no previous first episode psychosis study or study of people with prodromal symptoms of psychosis has been conducted. The purpose of the perceptual priors task was to measure precisely how much lip-movements influence what participants hear by using a staircase procedure (Cornsweet, 1962), in which the balance between two sounds was changed in predefined steps, providing a more fine-grained measures of individual susceptibility to the illusion than in previous clinical studies.

The second paradigm (from now on ‘cognitive priors task’), assesses the influence of learned written-word-sound associations on auditory perception. The impact of learned associations on auditory perception has been shown in sensory conditioning, where one stimulus functions as a predictor for an auditory stimulus that is otherwise difficult to detect. In these early experiments, participants were asked to identify auditory stimuli on the basis of a visual cue. Sometimes the participants reported perceiving an auditory stimulus when only presented with the visual cue, as the brain predicted an auditory stimulus on the basis of the cue (Ellson, 1941; Kot et al., 2002; Warburton et al., 1985;Agathon et al., 1973; Brogden et al., 1947; Powers et al., 2017). Previous research found that this omission effect is stronger in individuals with hallucinations (Kot et al., 2002; Powers et al., 2017), suggesting an increase in the influence of learned ‘cognitive’ expectation on auditory perception in psychosis, in contrast to the diminishment in the influence of ‘sensory’ expectations in schizophrenia discussed above. However, up to date, no study has explored the influence of learned cognitive expectations in individuals at-risk for psychosis and compared it to the influence of sensory expectations on perception.

We recognize that the sensory and cognitive priors tasks are strictly speaking not able to estimate the relative precision and mean of the prior expectations and sensory evidence for each participant directly. Instead we make the assumption based on Bayesian theories of the brain that perception is a function of the precision and mean of the prior and sensory evidence. Therefore rather then estimating the precision and mean for the prior and sensory evidence separately, we infer the relative contribution of prior information and sensory evidence, and term this for the remainder of this paper the relative strength of the sensory and cognitive prior. Reconciling the exact level of priors used in the current experiment in relation to the exact level of priors used in previous experiments in schizophrenia spectrum patients is not trivial. However, this is not central to our experiment. Our aim is to examine the effects of two different levels of priors on a given process at different stages of psychosis.

Another issue currently understudied relates to the neurobiological underpinnings of alterations in the contribution of prior expectations in perception. Changes in glutamate levels have been associated with schizophrenia (Marsman et al., 2011; Merritt et al., 2016; Treen et al., 2016), including in the cingulate cortex, where there is evidence of excessive glutamate in early illness stages, possibly progressing to reductions in later stages (Merritt et al 2016, Kumar et al 2018). It remains unclear to what extent glutamate levels in the brain relate to predictive coding mechanisms putatively mediating psychosis, in spite of various theoretical arguments and extrapolations from preclinical experiments (Corlett et al., 2011; Sterzer et al 2018). Notably the anterior cingulate cortex (ACC) has been associated with processing uncertainty (Rushworth et al., 2008) and precision-weighting of information in health and psychosis (Cassidy et al., 2017; Katthagen, et al., 2018; Haarsma et al., 2019). Thus alterations in glutamate levels in the ACC might alter the precision of prior information, thereby changing the degree to which priors influence perception. We therefore explored this issue by measuring magnetic resonance spectroscopy (MRS) glutamate levels in the anterior cingulate cortex and relating these measurements to the contribution of prior expectations in the different experimental groups. Our study is not powered to provide definitive results relating glutamate measures to our predictive coding measures, the latter being of primary interest here. Nevertheless, we report preliminary, exploratory analyses that may be hypothesis generating and could provide the basis for power calculations for future studies combining MRS with behavioural data in patients.

In summary, we use a cross-sectional design to study altered use of prior expectations in auditory perception in individuals at-risk for psychosis, first episode psychosis and controls. We expect to find differences in the balance between the use of prior expectations and sensory input depending on the origin of the prior expectation (sensory vs. cognitive) and disease stage (at-risk vs. first episode psychosis). Specifically, we expect that at early stages of psychosis (clinical risk), patients make relatively stronger use of sensory input then prior expectations relative to controls and individuals with a full manifestation of illness (first episode psychosis), but that in those with first episode psychosis, patients would rely more on cognitive priors relative to sensory input compared to controls and individuals at risk for psychosis. A secondary hypothesis is that cortical glutamate levels will be related to changes in the usage of sensory and cognitive priors.

## 1.2. Method

### 1.2.1. Participants

Participants with first episode psychosis (FEP, n=30, average 24.8 years, 6 female) or at-risk mental state patients (ARMS, n=29, average 21.5 years, 8 female) were recruited from the Cambridge Early intervention service North and South. In addition, ARMS patients were recruited from a help-seeking, low-mood, high schizotypy sub-group following a latent class analysis on the (Neuroscience in Psychiatry Network (NSPN) cohort (Davis et al., 2017) or through advertisement via posters displayed at the Cambridge University counselling services. Individuals with FEP or at-risk mental states for psychosis met FEP or ARMS criteria on the CAARMS interview. All FEP participants had current delusions or previous delusions in the case of those with partial or recent recovery. Healthy volunteers (Healthy control sample HCS, n=32, average 22.6 years, 15 female) without a history of psychiatric illness or brain injury were recruited as control subjects. Healthy volunteers did not report any personal or family history of neurological, psychiatric or medical disorders. All participants had normal hearing and normal or corrected to normal vision. All participants gave informed consent. The study was part of the NCAAPS study (Neuroscience Clinical Adolescent and Adult Psychiatry Study), which was approved by the West of Scotland (REC 3) ethical committee. See Table 1 for details on demographics and symptom scores. 3 ARMS patients and 17 FEP patients were receiving anti-psychotic medication.

**Table 1:** Demographics and symptom scores participants in the study

|  | HCS | ARMS | FEP | *p-value* |
| --- | --- | --- | --- | --- |
|  | 32 | 29 | 30 |  |
| **PANSS** | 13.1(4.6) | 26.7(12.1) | 31.6(12.3) | <.001 |
| *Positive* | 6.5(2.3) | 13.6(5.7) | 18.0(6.9) | <.001 |
| *Negative* | 6.6(2.4) | 13.1(7.5) | 13.6(7.7) | <.001 |
| **MFQ** | 8.5(5.1) | 33.2(17.4) | 31.8(23.6) | <.001 |
| **CAPS** | 32.9(1.4) | 44.1(7.0) | 43.6(9.7) | <.001 |
| *Distress* | 1.6(3.0) | 29.8(20.9) | 32.1(33.9) | <.001 |
| *Intrusive* | 2.2(3.7) | 34.9(22.8) | 38.5(37.4) | <.001 |
| *Frequency* | 1.3(2.3) | 28.3(17.8) | 29.7(31.1) | <.001 |
| **PDI total** | 22.4(1.5) | 29.3(4.5) | 31.1(6.5) | <.001 |
| *Distress* | 2.4(2.8) | 24.1(16.9) | 28.0(23.9) | <.001 |
| *Intrusive* | 2.4(2.7) | 23.6(17.4) | 29.5(22.9) | <.001 |
| *Conviction* | 3.6(4.0) | 24.9(15.9) | 31.0(25.3) | <.001 |
| **Age** | 22.4(3.7) | 21.8(3.5) | 25.1(4.8) | <.01 |
| **N Males** | 17 | 21 | 24 | >.05 |

### 1.2.2. Questionnaires and interviews

We used the Cardiff Abnormal Perceptions scale (CAPS, Bell et al., 2006), Peters Delusion Index scale (PDI, Peters et al., 1999), Comprehensive Assessment for the At-risk Mental State interview (CAARMS, Yung et al., 2003) and Positive and Negative Symptoms Scale (PANSS, Kay et al., 1989) to assess “caseness”, symptom severity and frequency. Both the total scores for the CAPS and PDI and the subscales of the CAPS and PDI are reported in table 1. For the PDI and CAPS the participants were required to give a yes or a no answer to a particular question. In case of a yes answer, 3 subscales were filled in which utilised a 5-point Likert scale. The CAARMS and PANSS are semi-structured interviews, where the interviewer rates severity of various types of psychotic and other psychiatric symptoms.

### 1.2.3. Magnetic Resonance Spectroscopy

A subset of participants was scanned on a Siemens Prisma 3T scanner at the cognition brain sciences unit in Cambridge. The spectroscopy scan was part of a larger MRI protocol which contained in addition 2 fMRI protocols and a structural scan totalling 90 minutes. The structural scan was used to plan the MRS voxel. A 15mm isotropic voxel was placed carefully in the anterior cingulate cortex. A PRESS sequence was used to assess glutamate levels, with a TR of 1880ms and TE of 30ms. 150 water-suppressed acquisitions were collected in addition to 16 unsuppressed acquisitions. Data was analysed in LCModel. MRS data was successfully collected from 18 healthy controls 19 ARMS, and 14 FEP patients.

### 1.2.4. Experiment 1-providing perceptual priors

In the present study auditory stimuli were presented that contained varying proportions of the phoneme /Ba or /Da (Figure 1). The balance between the two stimuli always adds up to one. The contribution of the stimulus /Ba is denoted as ω^Ba^, which stands for “the weight of /Ba”. The proportion of ω^Da^ can be derived from ω^Ba^ as 1 - ω^Ba^ = ω^Da^. From henceforth the notation ω^Ba^ be used to indicate what exactly was presented to participants in terms of auditory stimulus.

**Figure 1:**
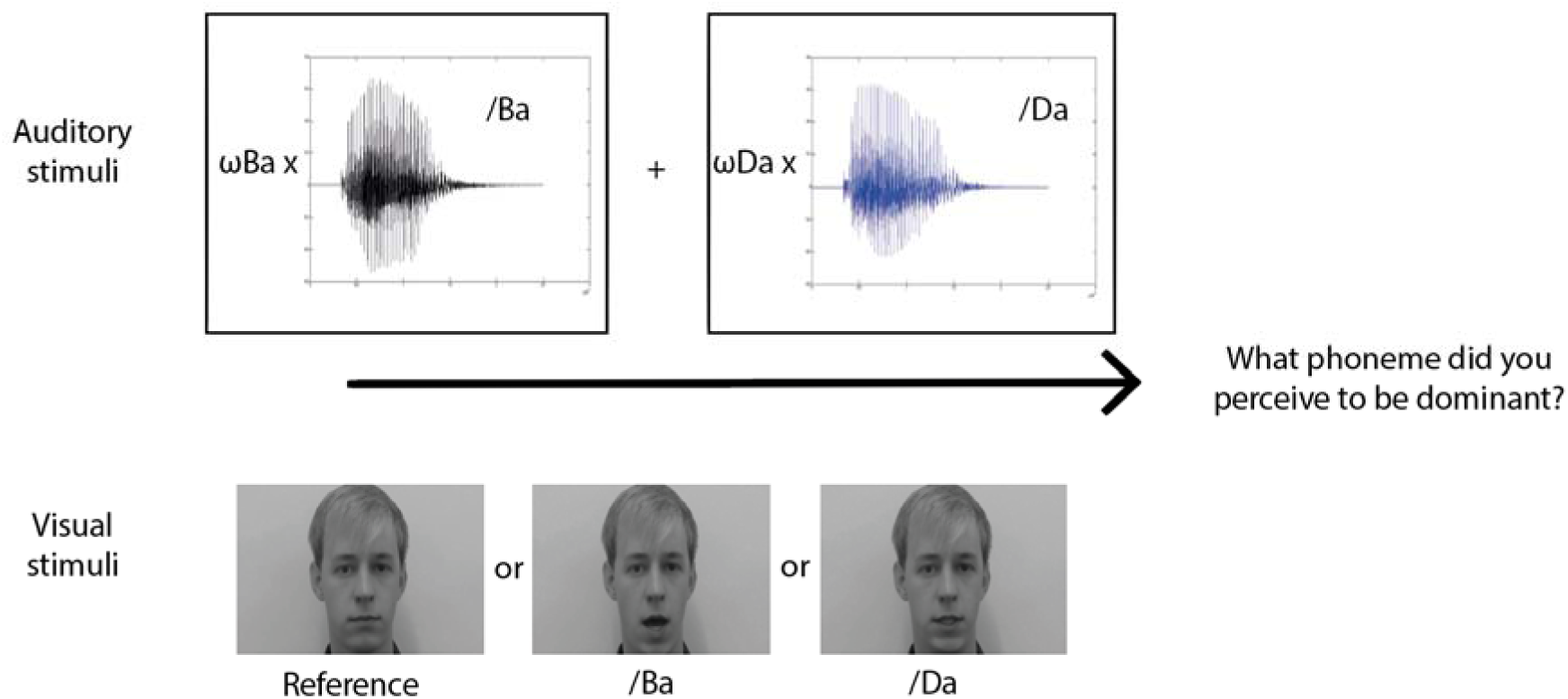
Procedure of the sensory prior task. The participant was presented between a mixture of the phonemes /Ba and /Da (above) which co-occurred with either a still face (reference condition) or lip-movements pronouncing /Ba or /Da

#### Training phase

The task started with a training phase. The purpose of which was to familiarize the participants with the auditory stimuli. Here they were presented with a still face in combination with an auditory stimulus consisting of a stimulus ω^Ba=^ .8 or ω^Ba=^ .2. They were then asked to report which sound they believed was dominant, after which they received feedback (correct/incorrect). The training was completed as soon as participants reported the correct answer 4 times for each stimulus. All participants identified the phonemes correctly.

#### Testing phase

During the testing phase, the participants were presented with an auditory stimulus consisting of a mix between the sound /Ba and /Da (as described above), which simultaneously occurred with a visual stimulus consisting of a black and white male face. The face would pronounce either /Ba or /Da (lip-movement condition), or the face would remain still (the reference condition). All three conditions were presented in a pseudo-randomised order such that all three conditions were presented in a random order before one of the conditions is presented again. The participants were instructed to keep looking at the lips of the face throughout the task, but asked to report what phoneme was dominant in the auditory stimulus by pressing one of four buttons indicating the level of certainty and the perceived phoneme.

During the main task, the balance between the /Ba and /Da phoneme was changed in a stepwise fashion. That is, when the participant reported the sound /Ba to be dominant in for example the reference condition, then the next time that condition came up, the balance between the sound /Ba and /Da would have been shifted in favour of the non-reported phoneme, in this case: /Da. By following this procedure, the task would converge towards a point where the participant would find it difficult to distinguish which of the phonemes is dominant in the auditory stimulus. This point is referred to as the perceptual indifference point. In the reference condition, where no lip-movements were presented, we expected the perceptual indifference point to converge on a stimulus which contains .5 of /Ba and .5 of /Da. However, when lip-movements, for example pronouncing /Ba were presented to bias perception towards the prior expectation, we expected that the task converged upon an indifference point that contained less auditory /Ba, and more auditory /Da. In other words, more auditory /Da was needed to overcome the influence that the /Ba lip-movements had (see Figure 2, top panel, for a schematic representation of the perceptual staircase experiment and figure 3 for an example of a staircase).

**Figure 2:**
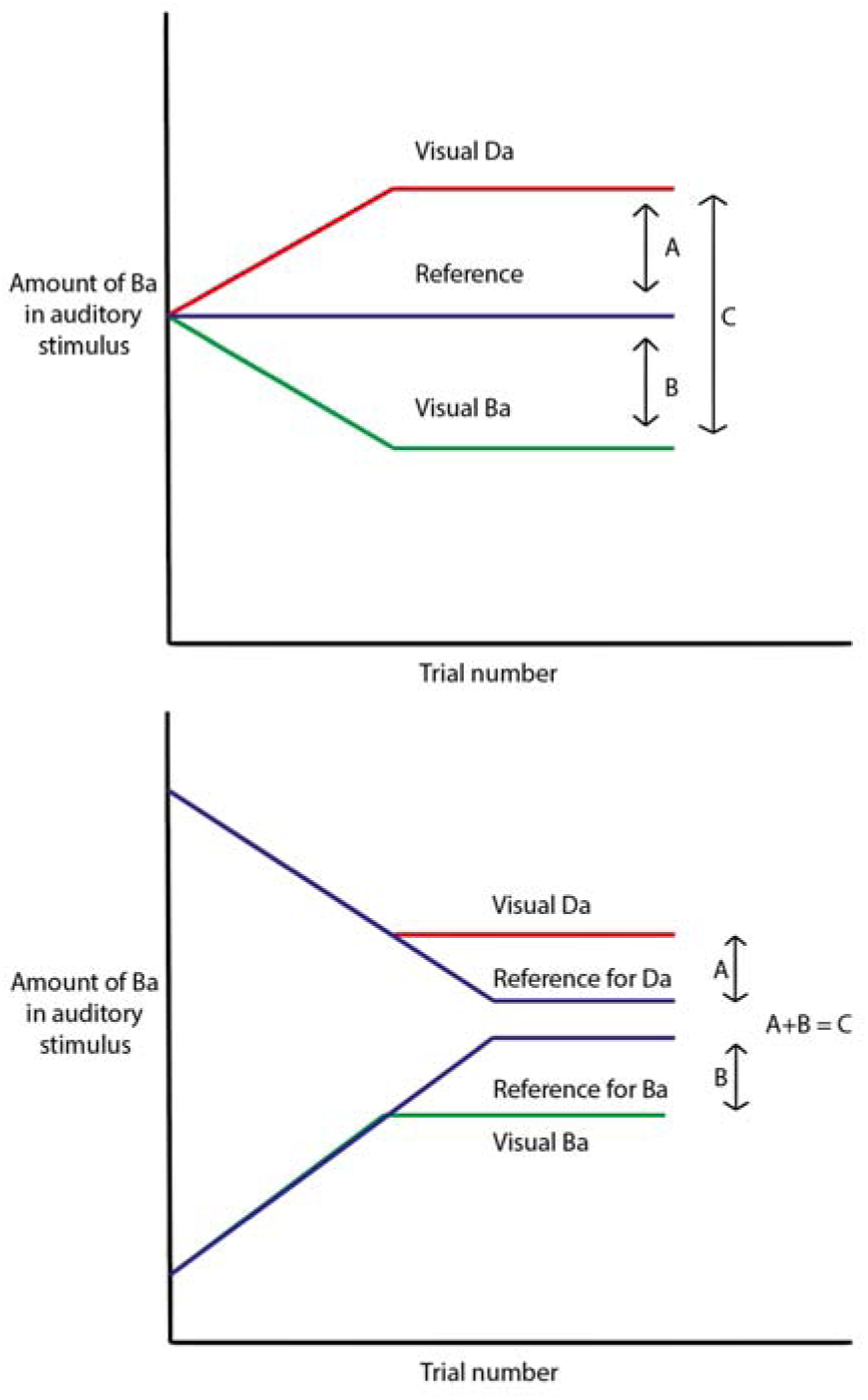
Schematic representation of a staircase in the perceptual priors task (upper panel) and cognitive priors task (lower). The experiment adjusted the balance between / Ba and /Da during the experiment in favaur of the nan-reported stimulus (slope line), ensuring convergence to a subject threshold (flat line). The distance A indicates the strength of the Da prior, whereas B indicates the strength the Ba prior. C is a total measure of prior strength irrespective of the specific priar presented.

**Figure 3:**
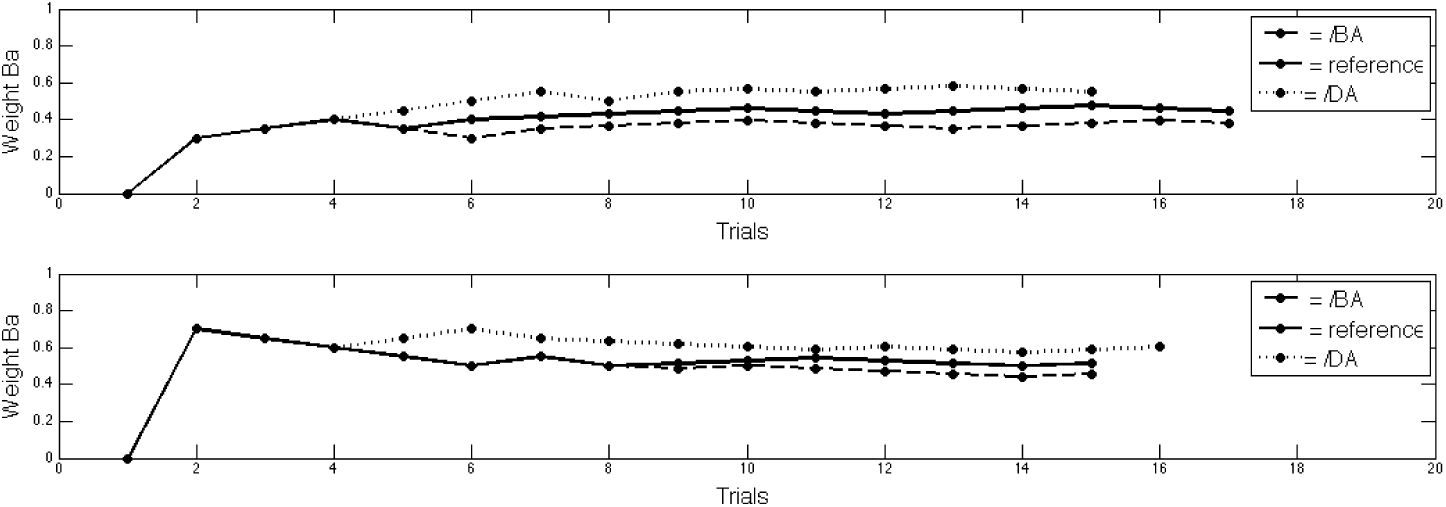
Example of the staircase procedure. All 6 of the conditions are represented here. The top figure shows the 3 visual conditions where the staircase started at ω^Ba^=.3, whereas the bottom figure shows the 3 visual conditions where the staircase started at ω^Ba^=.7.

For each of the three conditions (Reference, /Da and /Ba), the perceptual indifference point was assessed twice : Once where the auditory stimulus started with a dominant /Ba stimulus (ω^Ba=^ .7, ω^Da=^ .3) and once where /Da was dominant (ω^Ba=^ .3, ω^Da^= .7). This created 6 conditions, which were presented to the participant in pseudorandom order. A condition was completed when either one of two criteria was met. First, in the majority of cases, a perceptual indifference point was reached which was defined as having made 6 switches in perceiving one stimulus over the other (e.g. previously perceiving /Ba on trial t-1 and perceiving /Da on t0, indicating the balance between the two auditory stimuli is close to the participants perceptual indifference point). Second, a condition was completed when the participant indicated that the sound /Ba or /Da is 100% dominant in the auditory stimulus (e.g. a participant perceived /Da, even though the stimulus is 100% /Ba/ which could happen when the visual priors are dominating perception). In the second case this would technically not be an indifference point. However, for the remainder of this study we will refer to it as such for the sake of simplicity. The priors dominated perception only in a small minority of cases (see results). A condition was aborted when 30 trials had been presented avoiding the task from taking too long. This did not change the way the effect of the prior was calculated. In order to test for group and condition differences in the amount of trials needed to reach an indifference point and a possible interaction, we used a mixed-ANOVA with group as between subject factor and visual condition as within subject factor.

At the beginning of the staircase, the balance between /Ba and /Da was changed in steps of .05. After the first switch, the balance was changed in steps of .015. This procedure ensured that the first switch was reached quickly.Thereafter the staircase became more sensitive so that the perceptual indifference point could be assessed more precisely. The strength of each of the visual priors was calculated separately by taking the difference between the perceptual indifference point of the visual prior condition and the reference condition (see Figure 2 upper panel: A and B). The total strength of the visual priors was calculated by taking the distance between the indifference points of both sensory prior conditions (see Figure 2 upper panel: C).

### 1.2.5. Experiment 2 - providing cognitive priors

#### Training phase

The cognitive priors tasks was designed to measure how much a learned cue would influence what participants hear. During the training phase participants learned the association between the letters BA and the phoneme /Ba, and vice versa for DA. In 75% of the training trials the letters BA or DA were presented 500ms prior to hearing the auditory stimulus which consisted of ω^Ba=^.3 and ω^Da=^ .7 when preceded by the letters DA or ω^Ba=^ .7 and ω^Da=^ .3 when preceded by the letters BA, making the letters predictive of the auditory stimuli. In the other 25% of the trials, no sound was presented following the letters. Here the participants were asked to report what they expected to hear. The training was complete as soon as the participants indicated 8 times that they expected to hear the /Ba following the letters BA and /Da following the letters DA.

#### Testing phase

The cognitive priors task is similar to the perceptual priors task, in that participants were instructed to report which sound they believed to be dominant under different prior expectations. However, this time the prior expectations came from learned written word-sound associations. Again, the main task consisted of 3 conditions, a cognitive prior BA and DA condition, and a reference condition, which consisted of the letter ‘?A’. Each trial started with the presentation of the letters ‘BA’, ‘DA’ or ‘?A’. After seeing ‘BA’ or ‘DA’, participants were asked which phoneme they expected to perceive, which they indicated using one of 4 buttons indicating the perceived phoneme and certainty like in the perceptual priors task. The participants were only asked to indicate their prediction following seeing the letters ‘BA’ and ‘DA’, but not after seeing ‘?A’. By making a conscious prediction regarding the upcoming stimulus, the use of the cognitive prior could be validated. In the reference condition, no reliable prediction could be generated as both options were equally likely. 500ms after they made a decision or the reference stimulus had been presented, the auditory stimulus was presented. Subsequently, participants indicated what they perceived to be the dominant stimulus (see Figure 4).

**Figure 4:**
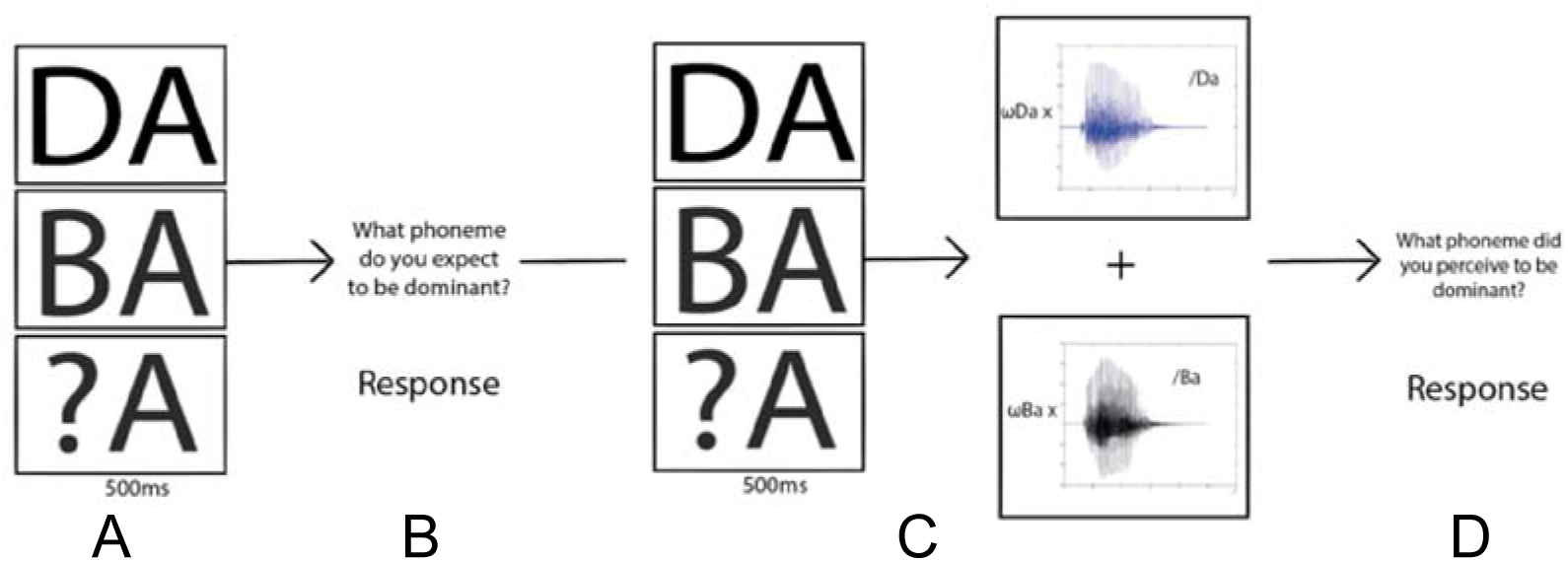
Procedure of the experimental phase of the cognitive prior task. A: First one of the three sets of letters was presented to the participant to indicate what sound was most likely to occur according to the training phase. B: participants were required to indicate which phoneme they believed to be most likely presented. C: The participant was again presented with one of the three letters (the same as in A) and 500ms later was presented with the mixed phoneme. D: After the presentation of the sound the stimuli were removed from the screen and the participant was required to indicate what phoneme they perceived to be dominant.

Again, the balance between the auditory phoneme /Ba and /Da was shifted in favour of the non-reported stimulus in a step-wise fashion. However, in contrast to the perceptual priors task, each condition was presented once for each cognitive prior BA and DA, instead of twice. Within the cognitive BA prior condition, the staircase started at ω^Ba^ = .7 and ω^Da^ = .3, meaning the auditory stimulus was relatively clearly a /Ba sound. The same is true for the cognitive DA prior condition, where the staircase started at ω^Ba^ = .3 ω^Da^ = .7, meaning the auditory stimulus was relatively clearly a /Da sound. This matching of the auditory stimulus to the cognitive prior condition at the beginning of the staircase was done to reaffirm the association between the prior and the sound, otherwise the association between the cue and sound could have been lost immediately in the beginning of the staircase. Note that if we would compare the difference in perceptual indifference points in the two cognitive prior conditions, we would have a confound, as the staircases for the two cognitive prior conditions started at different intensities, explaining any differences between the two conditions. Therefore, we introduced two reference conditions to which the prior conditions can be compared, getting rid of the confound. These consisted of the letters ‘?A’, one of which had a staircase starting at ω^Ba^ = .7 and ω^Da^= .3 so it could be directly compared to the cognitive BA prior, the other starting at ω^Ba^ = .3 ω^Da^ = .7, so it could be directly compared to the cognitive DA prior. As in the first task, at the beginning of the staircase procedure, the balance between /Ba and /Da was again changed in steps of .05. Then, after the first switch, the balance was changed in steps of .015.

In total, the cognitive priors task consisted of 4 conditions: a BA and a DA condition, a reference condition for BA, and a reference condition for DA. The order of the condition per participants was pseudorandomised. In each condition, a perceptual indifference point was assessed.

The perceptual indifference point for each condition was quantified by taking the average of ω^Ba^ at the last two switches. We also briefly rapport the results for taking the final four switches to demonstrate this does not influence the results substantially. In order to quantify the strength of each prior, these perceptual indifference points were subtracted from their reference condition, and the total cognitive prior strength was calculated by adding the strength of separate priors (see Figure 2 lower panel).

### 1.2.6. Stimuli, Apparatus and Procedure

Participants completed two tasks: the perceptual priors task first and the cognitive priors task second. Each task was performed on a MacBook Pro, Retina, 13-Inch, Early 2013, and each lasted on average about 10 minutes. Participants wore Sennheisser Headphones to ensure optimal hearing. Both the Ba and the Da stimuli had an intensity of 68dB. All participants reported perceiving the auditory stimuli clearly. The experiment was conducted in an environment with minimal background noise, ensuring minimal distraction of the participant (<15dB).

Psychtoolbox-3 was used to design the experiment. The auditory stimulus in both the perceptual priors task and the cognitive priors task consisted of a mixture of a natural speech male voice /Ba phoneme and a /Da phoneme. The auditory stimulus was created by multiplying the auditory spectrum of the /Ba stimulus by a weighting factor ω^Ba^. This was then added to a weighted auditory spectrum of /Da (where ω^Da^=1-ω^Ba^) ensuring the total of auditory stimulus to always be 1(stimulus = (ω^Ba^ x Ba) + (ω^Da^ x Da)).

### 1.2.7. Analyses

Since this is a novel paradigm, we first wanted to establish whether the variables of interest were reliable in the sense that two separate measurements of the variable were highly correlated. Since we assessed the perceptual indifference points twice in each condition, we were able to test the correlation between two separately obtained measurements, giving an indication of their reliability. We tested the reliability of two separate variables. First, we tested the reliability of the indifference points in the condition without a perceptual prior, which should give an indication of the reliability of the staircase method. Second, we tested the reliability of the strength of the perceptual priors, which give an indication of the reliability of the method to measure the influence of lip-movements on auditory perception. Furthermore we tested whether the perceptual and cognitive priors were correlated with each other. Due to non-normality of the cognitive priors task, a Spearman correlation was used to assess this.

One tailed paired T tests were used to test for a main effect of whether the lip-movements shifted the perceptual indifference points in the expected direction compared to the reference condition. This was done for both the sensory and cognitive prior tasks.

In order to test the hypothesis that perceptual priors and cognitive priors were different across groups, we computed the influence of the prior for each individual as described above, and used a one-way ANOVA with two-tailed post-hoc Bonferroni corrected t-tests if applicable. Furthermore, a Kruskal Wallis non-parametric ANOVA was used with cognitive prior data, with Bonferroni corrected non-parametric post-hoc t-tests. We also report the results of Bayesian statistical tests in relation to the group differences using JASP. We report effect sizes for the key statistical tests, i.e. effect of group on prior strength. We report Cohen’s d for T-tests, and *η*^2^ for the one-way ANOVA’s. All effect sizes are calculated on the basis of parametric tests.

## 1.3. Results

### 1.3.1. Perceptual priors task

#### 1.3.1.1. No difference between groups in the amount of trials needed to assess perceptual indifference point

On average participants required 18.9 trials to reach a perceptual indifference point across all conditions. We found no overall effect of group on the trials needed to reach a perceptual indifference point (F{2,87}= .262, *p*=.77) (HCS: 19.1, SE: 0.5; ARMS: 19.1, SE:0.6; FEP: 18.6, SE: 0.4). However, we did find an effect of prior condition (F{2,174}=17.1, *p*<.001): needing fewer trials in the visual reference condition (17.3, SE: 0.3) than in the visual BA (18.9 SE: 0.4) and visual DA condition (20.7, SE: 0.54). Importantly, we found no group by condition interaction (F{4,174}=.456, *p*=.77). Thus, the patient groups did not differ in terms of the trials needed to reach indifference points.

#### 1.3.1.2. Individual perceptual indifference points can be estimated reliably

The perceptual indifference point for each visual condition was assessed twice in the perceptual priors task. As this is a novel task, we tested whether these simultaneously assessed indifference points correlated strongly, as that would give us an indication of the reliability of the measurement. First, we correlated the indifference points in the condition where no priors were presented (the reference condition). Across groups the correlation was r=.73. Separately it was r=.83 for HCS, r=.76 for ARMS and r=.55 for FEP (all *p*<.01). The correlation between the two reference points was significantly higher in the HCS group compared to the FEP group (Fisher r-to-z transformation: *p=* .033), but not between other groups all *p>* .2. Second, in a similar fashion, we assessed how strongly the effect of the perceptual priors was correlated across the two simultaneously assessed staircases. The reliability of the strength of the perceptual priors across groups was r=.78. Separately, it was r=.88 for HCS, r=.79 for ARMS and .69 for FEP (all *p*<.01) (Figure 5). The differences in correlations between perceptual priors were not significantly different *p*>.2. For the remainder of the analyses we averaged for each visual condition (Ba Da and reference) the perceptual indifference points, and the estimation of the sensory prior strength (Figure 2).

**Figure 5:**
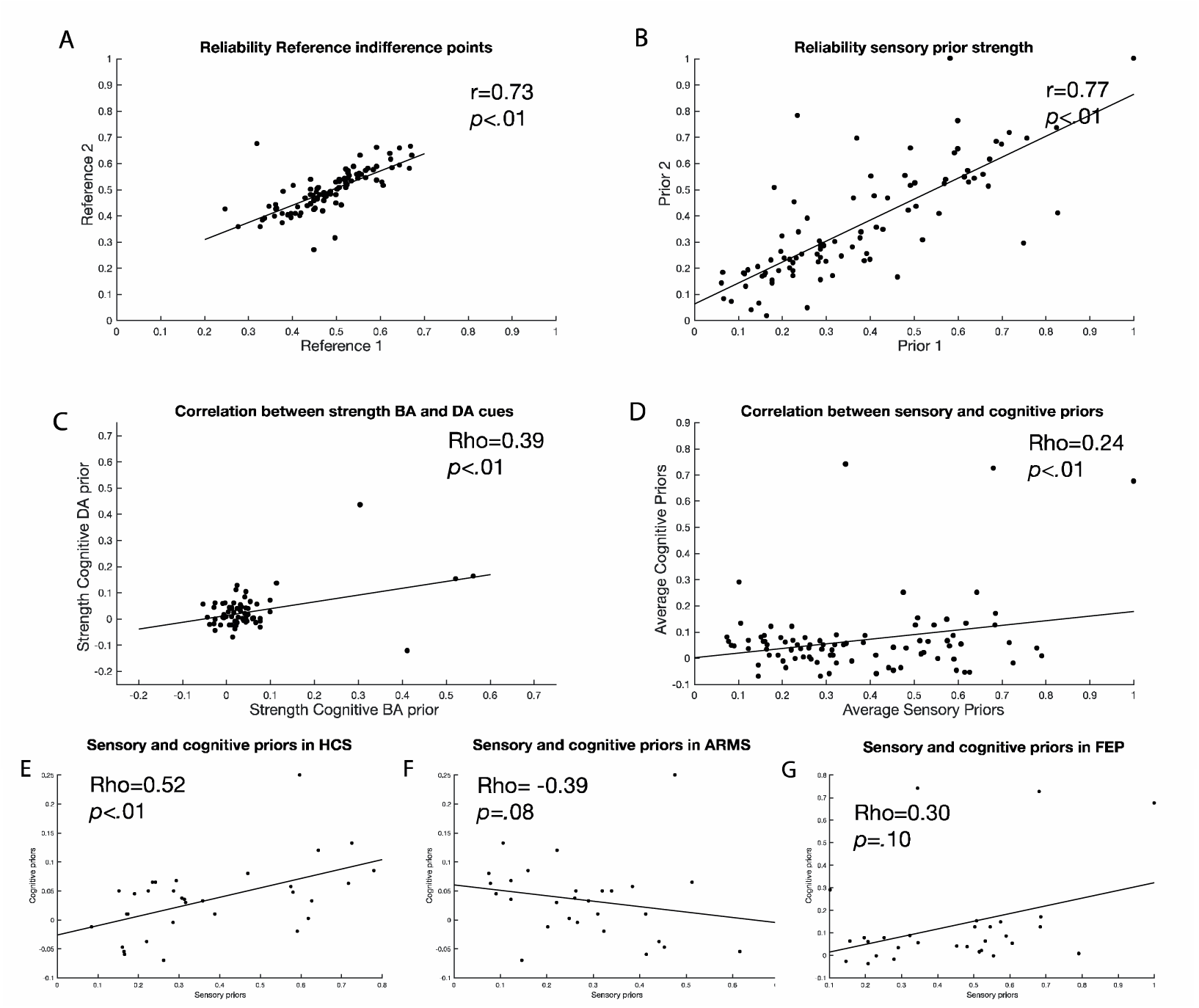
Correlations testing the reliability of the experiment are presented here. A: reliability of the perceptual indifference point in the reference condition. B : reliability of the strength of perceptual priors. C: Correlation between the effect of cognitive Ba stimulus and the cognitive Da stimulus. D: correlation between sensory and cognitive priors. E-G: relationship between cognitive and sensory priors for each experimental group. Whereas healthy controls and FEP show a positive correlation, ARMS shows a negative correlation. We calculate Spearman correlations but include linear fit lines for display purposes.

#### 1.3.1.3. Perceptual priors shifted the perceptual indifference points in the expected direction

We tested whether the perceptual priors shift the perceptual indifference points in the expected directions compared to the reference condition. On average, across all groups taken together, Ba lip-movements lowered the value of ω^Ba^ in the perceptual indifference point by .21 (95% ci: .18-.23, T{89}=14.0, *p*<.0001). In contrast, Da lip-movements increased the value of ω^Ba^ in the perceptual indifference point by .16 (95% ci: .14- .18, T{89}=13.2, *p*<.0001) on average. When comparing the relative strength of the Ba and Da lip-movements, we found a significant difference (T{178}=2.29, *p*=.022), indicating a slightly stronger effect of Ba lip-movements then Da (Figure 6A).

**Figure 6:**
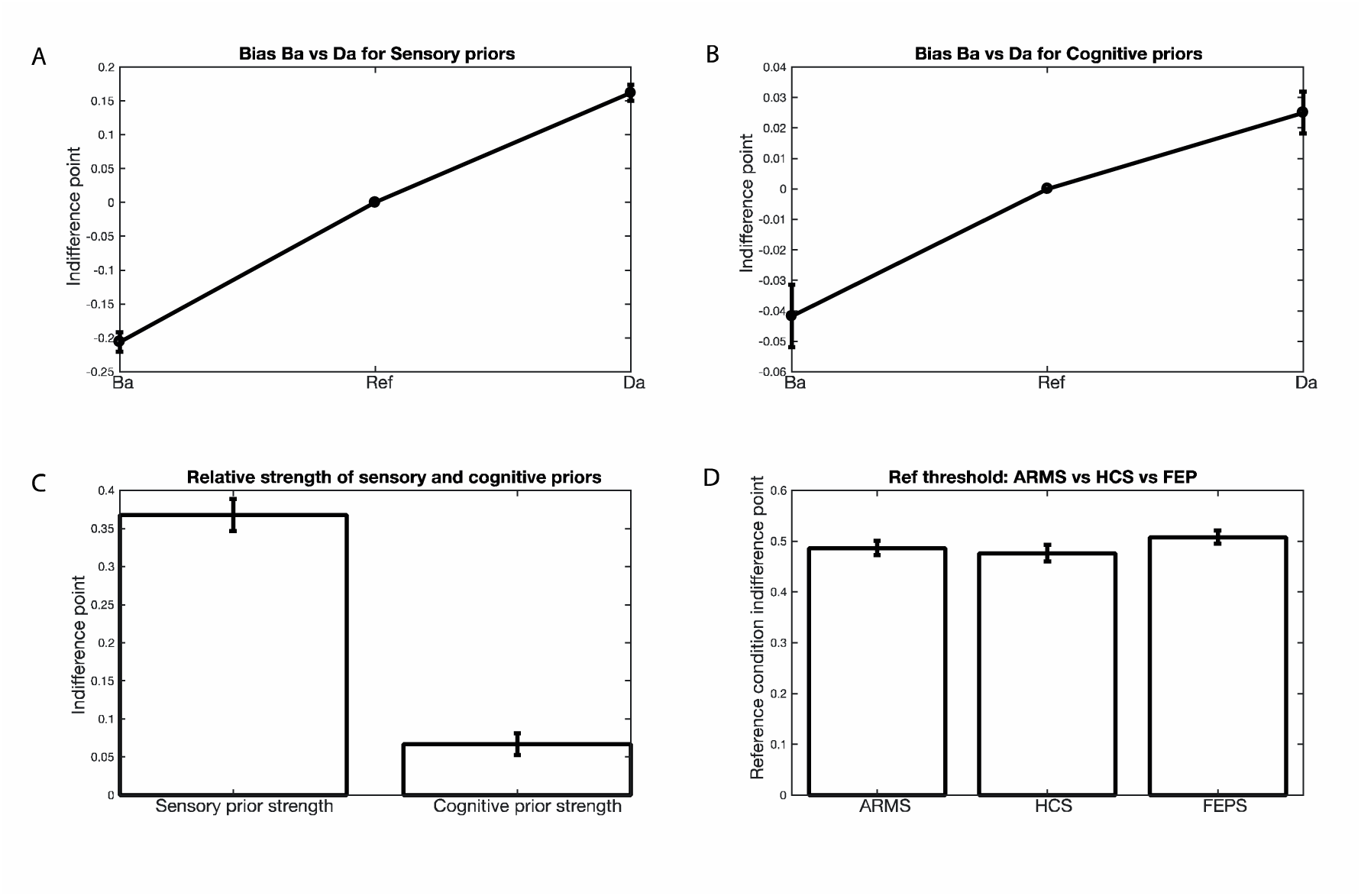
Main effects of the sensory and cognitive priors are presented here. A: relative shift in perceptual indifference points under different sensory prior conditions (lip movements pronouncing /Ba or /Da) compared to reference condition (still lips). B: relative shift in perceptual indifference points under different cognitive prior conditions (the letters ‘BA’ and ‘DA’) compared to reference condition (letters ‘?A’). C: relative strength of perceptual priors and cognitive priors. D: the perceptual indifference points in the reference conditions per group (effect of no interest). Error bars represent standard error of the mean.

#### 1.3.1.4. The perceptual indifference point in the reference condition was equal across groups

Analysing group differences, the perceptual indifference point in the reference condition was a variable of no interest, as it merely reflects a personal preference for either the auditory /Ba or /Da stimulus. Indeed, the average perceptual indifference point in the reference condition across groups in reference groups was equal (M^HCS^= .48 S^EHCS^= .02, M^ARMS^= .49 SE^ARM^=.01, M^FEP^=.51 SE^FEP^=.01; F{2,88}=1.02, *p*=.36) (Figure 6D).

#### 1.3.1.5. Perceptual priors were significantly lower in ARMS compared to FEP

To test whether the perceptual priors were significantly different across groups, we conducted a one-way ANOVA. We indeed found evidence for a difference across groups (F{2,88) = 5.32, *p=.007,* effect size *η*^2^ =.11; Figure 7A, 7C). Bonferroni corrected post-hoc T-tests revealed a significant difference between ARMS (M^ARMS^=.28 SE^ARMS^=.03) and FEP (M^FEP^=.44 SE^FEP^=.04) *(p=.005,* effect size d=.89, ci= .46-1.32), but not between healthy controls (M^HCS^=.37 SE^HCS^=.04) and ARMS (*p*=.20, effect size d= −.51, ci=.01-1.01) or FEP *(p=.44,* effect size d=.34, ci=-.13-.85). We tested whether changing the amount of switch points that were used to calculate the indifference point changed the results. When we change this from two to four, we find the same (slightly stronger) effect: F(2,88)=5.72, *p*=.005, ARMS vs FEP: *p*=.002, ARMS vs HCS: *p*=.12, HCS vs FEP: *p*=.24).

**Figure 7:**
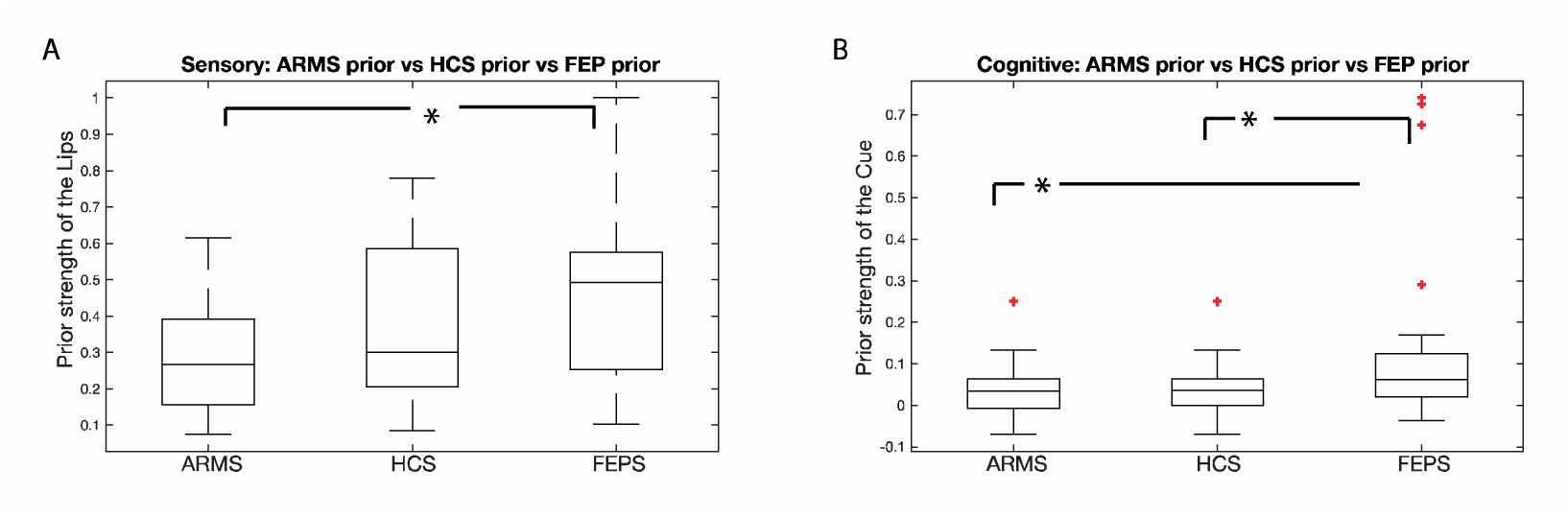
The effects per group are presented here in boxplot A: The effect of perceptual priors across groups. B: The effect of cognitive priors across groups. * = *p*<.05

We furthermore analysed the perceptual prior data in a Bayesian fashion. For this section we use Jeffreys’s (1961) suggested evidence categories for the Bayes factor. We found that an ANOVA revealed moderate evidence in support for a difference across groups (BF=6.3). Independent-sample t-tests revealed anecdotal evidence in favour of a difference between ARMS and healthy controls (BF=1.4), but anecdotal evidence in favour of no difference between healthy controls and FEP (BF=1.8). There was strong evidence for a difference between ARMS and FEP (BF=26.1) (Figure 7A, 7C).

The /Ba perceptual prior dominated perception completely in 4/32 HCS, 0/29 ARMS and 5/31 FEP participants, whereas the /Da perceptual prior dominated perception completely in 5/32 HCS, 2/29 ARMS and 11/31 FEP. In one FEP participant the both the /Da and /Ba lip-movements completely dominated perception.

### 1.3.2. Cognitive priors task

#### 1.3.2.1. FEP needed on average an extra trial to finish the training phase

We first tested whether the different experimental groups differed in the amount of trials needed to end the training using an ANOVA. The groups differed significantly in the number of trials needed (F{2,88}=3 .34, *p*=.040). The HCS group and the ARMS group required on average 8.7 trials and 8.8 trials respectively before the training was finished, whereas the FEP required on average 9.9 trials.

#### 1.3.2.2. No difference between groups in the amount of trials needed to assess perceptual indifference point

During the actual experiment, the participants generally required 18.5 trials to reach a perceptual indifference point across all conditions. We found no overall effect of group on the trials needed to reach a perceptual indifference point (F{2,88}=.44, *p*=.64) (HCS: 18.5, SE: 0.6; ARMS : 18.9, SE:0.6; FEP: 18.1, SE: 0.6). However, we did find an effect of prior condition (F{2,88}=3.56, *p*=.033). Needing significantly fewer trials in the DA condition (17.6, SE: 0.5) then in the visual BA (19.5 SE: 0.5) (T {180}=2.63, *p=.* 018) but not the reference condition (18.3, SE: 0.5) (T{180}=1.08, *p*=.56, Bonferroni corrected). Importantly, we found no group by condition interaction (F{4,176}=.27, *p*=.90). Thus, the patient groups did not differ in terms of the trials needed to reach indifference points.

#### 1.3.2.3. Cognitive priors shifted the perceptual indifference points in the expected direction

In order to assess the main effect of cognitive priors, each perceptual indifference point of the two cognitive prior conditions was subtracted from its own reference condition. We found that the cognitive BA prior lowered the value of ω^Ba^ by .042 (zval = −5.2, *p<* .0001), and for the cognitive DA prior the value of ω^Ba^ was increased by .027 (zval = 3.7, *p=* .0002). This shows that there was indeed a main effect of cognitive priors on perceptual indifference points. The relationship between effect of BA and DA priors is shown in Figure 5C. For the remainder of the analyses, the degree of influence of the BA and DA cognitive priors were added together and averaged in order to create a single measure of cognitive prior strength (see Figure 6B).

#### 1.3.2.4. Effect of cognitive priors in the FEP group was significantly higher than the ARMS and controls

We used a non-parametric ANOVA that is robust against Type I errors in non-normally distributed data. The differences between the average strength of the cognitive priors was significant (Independent-Samples Kruskal-Wallis Test: *p=.0 23,* effect size *η*^2^ =.11). Using a post-hoc Bonferroni corrected Wilcoxon rank sum test, we found stronger usage of cognitive priors in the FEP group compared to both the HCS group (zval: 2.35, ranksum: 840, *p=.037,* effect size d=.64, ci=.11-1.17), and the ARMS group (zval:2.35, ranksum: 714, *p=.037,* effect size d=.62, ci=.10-1.14), but between the HCS group and the ARMS group *p>.5.* We tested whether changing the amount of switch points that were used to calculate the indifference point changed the results. When we change this from two to four, we find the same (slightly stronger) effect: FEP vs HCS: *p=.015,* FEP vs ARMS: *p=.016,* HCS vs ARMS: *p*>.5).

We also analysed the cognitive prior data in a Bayesian fashion, and found that an ANOVA revealed moderate evidence in support for a difference across groups (BF=7.5). Independent-sample t-tests revealed moderate evidence in favour of no difference between ARMS and healthy controls (BF=3.5), but moderate evidence in favour of a difference between healthy controls and FEP (BF=3.5). There was also anecdotal evidence for a difference between ARMS and FEP (BF=2.8) (Figure 7B, 7D). Although we had no evidence that the extreme values represent measurement error, we analysed the results having excluded outliers in all three experimental groups (1 HCS, 1ARMS, 3 FEP). We found similar results (two sample t-test adjusted for multiple comparisons: averaging over final 2 switch points: HCS vs FEP: *p= .035,* ARMS vs FEP: *p=.038.* Final 4 switch points: HCS vs FEP *p=* .050, ARMS vs FEP *p=* .051).

For the cognitive prior experiment there was one FEP participant for whom the BA prior completely dominated perception, and 2 other FEP participants for whom the DA prior completely dominated perception, with no occurrences in ARMS or HCS. There were no participants for whom both the BA and DA cue completely dominated perception.

### 1.3.3. Perceptual priors had a stronger effect on perception than cognitive priors and were differently correlated across groups

Finally, we analysed whether the strength of the priors was different between tasks. This was indeed the case, showing a stronger effect of perceptual priors (.37) compared to the cognitive priors across all groups (.07) (T{90}=-14.34, *p<.0001,* effect size d=1.5, ci= 1.8-1.2) (Figures 5D, 6C). Subsequently, we tested whether the strength of cognitive and perceptual priors was correlated using a Spearman correlation. This was indeed the case (Rho=.24, *p*<.02). When exploring the correlations separate for each group, we found a negative (trend-level) correlation in the ARMS group (Rho=-.33, *p*=.08), and positive correlations in the HCS (Rho=.52, *p*=.002) and (trend-level) in the FEP group (Rho=.30, *p*=.10). Using a Fisher r-to-z transformation We found that the relationship was significantly different for the ARMS group compared to the healthy control group (Z=-3.28, *p*=.001), and FEP group (Z=-2.25, *p*=.024). The correlation between healthy controls and FEP was not significantly different (Z=1.0, *p*=.31). As these findings constituted secondary analyses, they are not properly controlled for multiple comparisons. When controlling for multiple tests, only the relationship in the healthy control group remains significant.

### 1.3.4. Glutamate levels correlate with cognitive priors in HCS and perceptual priors in FEP

Correlations with glutamate were tested in a subset of participants, namely 18 healthy controls, 19 ARMS, and 14 FEP patients. We looked for a correlation across all participants between glutamate levels and the strength of the perceptual and cognitive priors, but found no significant correlation (perceptual: Rho=.18, *p*=.21, cognitive: Rho=.17, *p*=.23). When exploring the correlations in the separate patient groups, we found that there is a significant positive relationship between glutamate levels and cognitive priors in the control group (Rho=.53, *p*=.023), but not with perceptual priors (Rho=.294, *p*=.24). In the ARMS group no significant correlations were found for either cognitive (Rho=.0, *p*=1) or perceptual priors (Rho=.07, *p*=.78). In the FEP group a significant correlation was found with perceptual (Rho=.57, *p*=.035) but not cognitive priors (Rho=.43, *p*=.128). As these findings were secondary to the core hypothesis in the present chapter, they were not corrected for multiple comparisons. The effects do not remain significant when they are controlled for multiple comparisons (See Figures 8 and 9).

**Figure 8:**
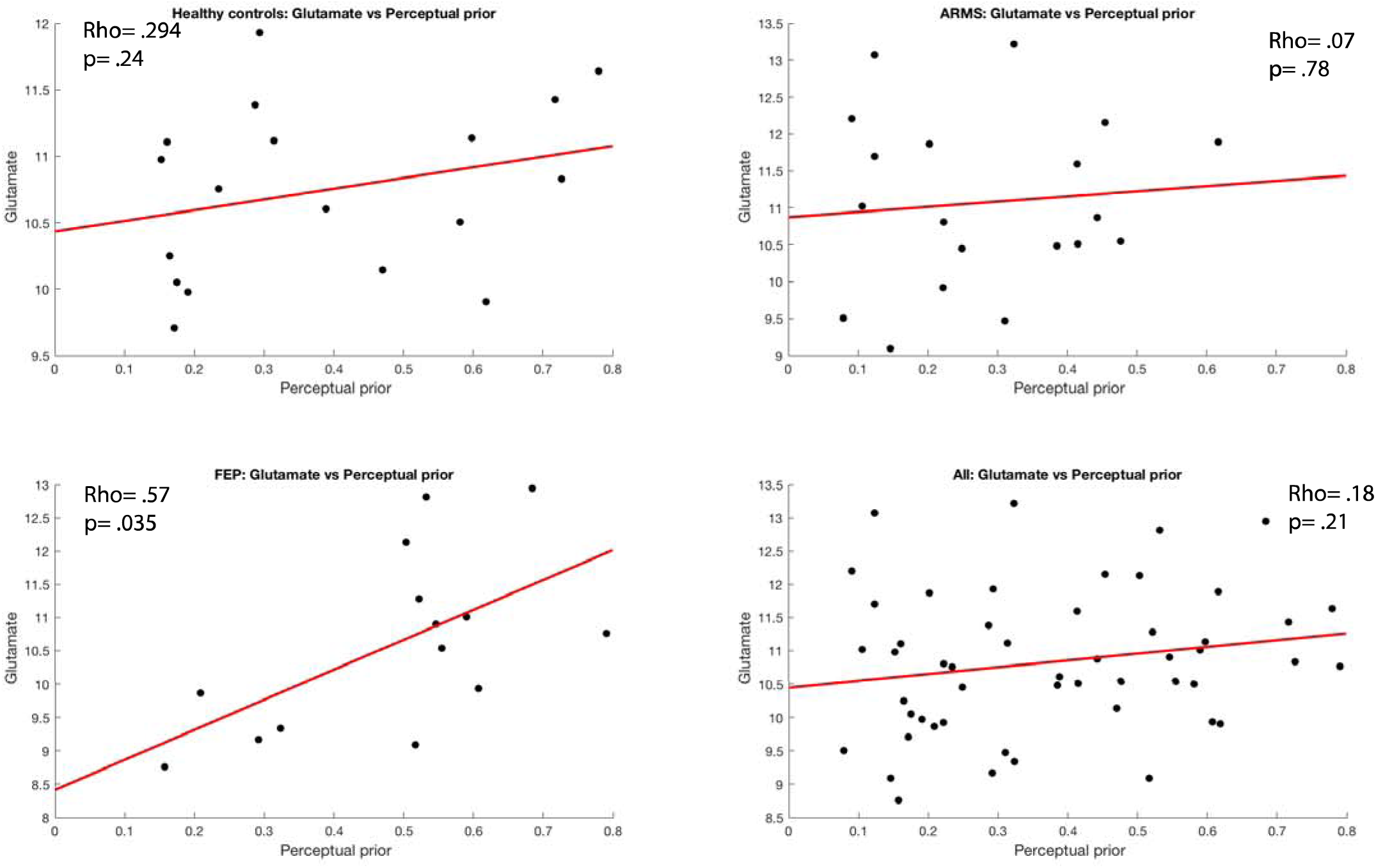
Correlations between perceptual prior strength and glutamate levels for all groups. We report Spearman’s correlations but plot linear fits for display purposes.

**Figure 9:**
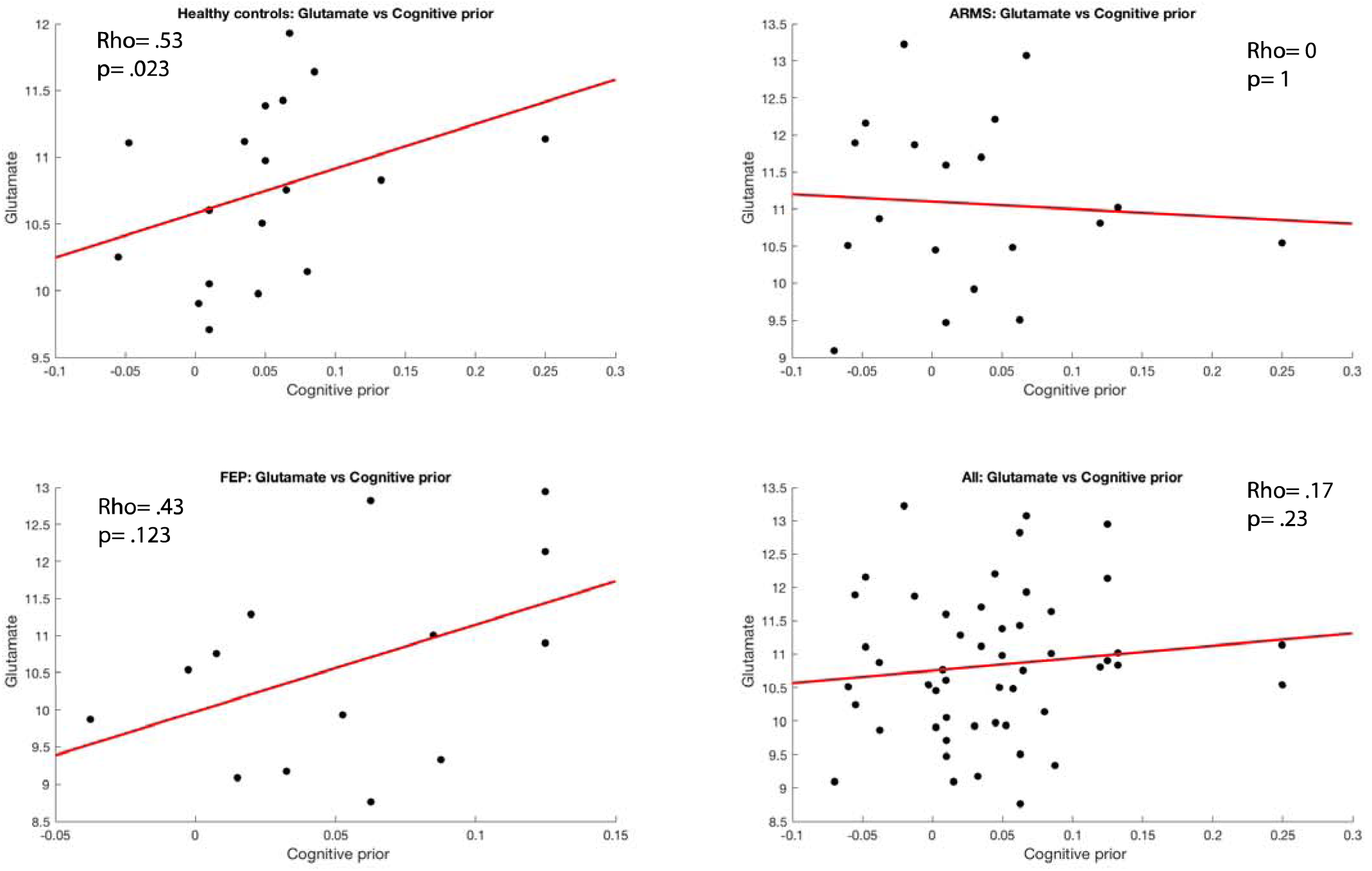
Correlations between cognitive prior strength and glutamate levels for all groups. We report Spearman’s correlations but plot linear fits for display purposes.

### 1.3.5. Stronger cognitive priors are associated with delusion ideation in ARMS and weaker perceptual priors is associated with delusion ideation and hallucinations in FEP

To explore the relationship between the usage of sensory and cognitive priors and the relation with symptoms, we computed Spearmen correlations within the different experimental groups (Table 2).

**Table 2:** Correlations between abnormal perception and belief and usage if sensory and cognitive priors across all groups.

|  | ARMS |  | FEP |  | ARMS+FEP |  | HCS |  |
| --- | --- | --- | --- | --- | --- | --- | --- | --- |
|  | Sensory | Cognitive | Sensory | Cognitive | Sensory | Cognitive | Sensory | Cognitive |
| PDI | p=.44<br>Rho=-.16 | p=.030<br>Rho=.44 | p=.023<br>Rho=-.48 | p=.19<br>Rho=-.29 | p=.09<br>Rho=-.25 | p=.28<br>Rho=-.16 | p=.92<br>Rho=.018 | p=.27<br>Rho=.21 |
| CAPS | p=.84<br>Rho=.044 | p=.87<br>Rho=.037 | p=.008<br>Rho=-.55 | P=.16<br>Rho=-.31 | p=.10<br>Rho=-.24 | p=.86<br>Rho=.03 | p=.06<br>Rho=.34 | p=.56<br>Rho=.11 |

In brief, we found that an increase in cognitive prior use was associated with delusion ideation in ARMS, whereas a decrease in the usage of perceptual priors was associated with perceptual abnormalities and delusion ideation in the FEP group.

## 1.4. Discussion

In the present study we found that whether prior expectations have a stronger or weaker effect on perception in psychosis depends on the origin of the prior expectation and the disease stage. We found strong evidence of weakened perceptual priors in the ARMS group compared to the FEP group, and some evidence of ARMS versus controls differences. In contrast, when comparing cognitive priors we found that the FEP group had stronger priors compared to the ARMS and healthy control group, whereas the healthy controls and ARMS group did not differ from each other.

The present findings can be interpreted in the hierarchical predictive coding framework. This framework suggests that the brain models the world by making predictions about upcoming sensory input, that are subsequently updated by discrepancies between the predictions regarding the sensory input and the actual sensory input, termed the prediction error (Knill et al., 2004; Friston, et al., 2005 & 2009; Rao et al., 1999; Bastos et al., 2012; Clark et al., 2013; Hohwy, 2014). In these models, abnormal perception and delusional beliefs can be expected to occur when the balance between the prior expectations and sensory input is shifted (Fletcher & Frith, 2009), as was found in the present experiment. That is, sensory input can come to dominate perception, likely resulting in the subjective experience of being overwhelmed by their sensory environment and attributing importance to otherwise irrelevant stimuli, as is sometimes reported in the early, including prodromal, stages of psychosis (Corlett et al., 2010, McGhie and Chapman, 1961; Bowers and Freedman, 1966; Freedman, 1974; Matussek, 1952).

Our results of abnormally strong high-level priors in first episode psychosis, all of whom had either current or recent delusions, are in accordance with previous postulates (e.g. Adams et al 2013, Sterzer et al 2018). We further note that high level, cognitive, priors were stronger in established psychosis compared to the ARMS, consistent with previous theory that strong high-level priors may develop subsequent to weak low-level priors (Adams et al 2013, Sterzer et al 2018). As Heinz et al (2018) reason, “reduced precision of *perceptual beliefs* encoded at low levels, e.g. in sensory cortices, may be compensated by increased precision of more abstract *conceptual beliefs* encoded in higher-level brain circuits.” However, previous theories have not described on what time scale this compensation happens, and no previous studies have examined over what time scale or at what stages in psychotic illness this may occur. Our data suggest that this compensation may not necessarily be instant, but might develop over time, possibly in the transition from the prodromal stage (ARMS) to frank psychosis (FEP).

A recent study examining the influence of prior expectations on auditory perception used a conditioning paradigm to study aberrancies in healthy voice hearers, voice hearers with psychotic illness, and psychotic illness without voice hearing (Powers et al., 2017). Individuals who heard voices were susceptible to report hearing a sound when none was present following a previously associated cue. Computational modelling showed that individuals with psychotic illness had difficulties learning that a cue failed to predict a sound, sticking to their prior expectations, whereas individuals who heard voices but did not have psychotic illness did recognise volatility and were able to alter high-level beliefs. This might in part explain why we only see an effect of the cognitive priors in the psychosis group, but not in the at-risk mental state group, who, although help-seeking, do not (yet) have psychotic illness. Because the current paradigm involves a staircase experiment, we only pick up strong effects of prior expectations in individuals who remain influenced by the priors towards the end of the experiment. The individuals at-risk for psychosis might have been influenced in the task early on, but changed their expectations regarding the validity of the cue later on. Since our key-variable is the influence of the priors at convergence, we might have been unable to pick this up.

It should be pointed out that there are a number of outliers in the first episode psychosis group. Although our statistical tests are robust against Type I errors in a data set with outliers (Zimmerman, 1994), and the results hold when removing these outliers, it still raises the question what the nature of the outliers is. One possibility is that there is a subset of individuals that is exceptionally strongly influenced by prior expectations. Indeed previous studies have reported non-normal data on such variables (see Powers et al., 2017 Fig 1E). However, there is also the possibility that these participants performed the task differently or misunderstood the instructions, although we have no evidence that these outliers were caused by experimental measurement error. The reliability of the perceptual priors was slightly less in the FEP group compared to the other groups, but it should be noted that this difference was not significant, and there was still a reliable correlation between the independently assessed prior strengths (correlation 0.7 for use of sensory priors in FEP). In addition an average was taken from the two independently assessed priors, likely increasing the reliability further. Furthermore, since the present task does not measure performance, but rather a perceptual bias, an increase in noisy decision making will not bias the results in one way or the other.

It has been argued that there might be a relationship between early sensory processing deficits and high level deficits in schizophrenia (Leitman et al., 2009). This raises the question what the exact nature of this exact relationship is and whether it might be relevant in understanding the development of psychosis. Whilst in the sample as a whole, cognitive and perceptual prior strengths were weakly (rho=0.24), but significantly, correlated, the strengths of the correlations were significantly different across groups. Although we acknowledge the caveat that within group correlations were of secondary interest in this study, and not well powered, the fact that the group comparisons in strengths of priors were sensitive to whether priors were high or low level provides supporting evidence that level of priors does matter in this research context. Perceptual priors in ARMS were negatively correlated with cognitive priors, whereas in FEP and healthy controls, and the sample as a whole, the correlation was positive. We speculate on the possibility that an increase in the influence of cognitive priors on perception in the FEP group is an adaptation to early visual processing deficits in the earlier stages of psychosis as seen in the at-risk group. This increase in cognitive priors subsequently could potentially act to counter the decrease in diminishment of perceptual priors explaining the positive correlation that is observed in the FEP group. This increase in cognitive priors may manifest themselves as delusions on the phenomenological level as can be seen in both the strong cognitive priors in FEP, and in the correlation with symptom severity in the ARMS group. Subsequently, if perceptual priors remain low in the FEP stage, this is correlated to worse symptomology, suggesting a failure for the brain to deal with a change in the perceptual system may be important for psychopathology severity in this stage of the illness. Interestingly, in the FEP stage there is no correlation between cognitive priors and symptoms, possibly due to noise added to the data through the effects of treatment, recovery in some, and delusional belief formation being an attempt at making sense of a changing sensory world (Mishara & Corlett, 2009). Overall, our data emphasize the importance of distinguishing between priors at high and low levels of the cognitive hierarchy (Schmack et al 2013).

We conclude that the initial stages of psychosis may be characterised by a weakening of lower-level perceptual priors. Compensatory neural systems changes may lead to deploying stronger higher-level priors in order to deal with the increased strong drive on perceptual input. These changes might be associated with formation of delusional beliefs (as supported by the correlations with symptoms). If this compensatory strategy is effective, the weakened perceptual priors may be restored throughout development. If ineffective, the perceptual priors remain weak and psychotic symptoms maintain (as supported by the correlations with symptoms). This model is described in Figure 10, where in red and blue the strength of perceptual and cognitive priors are depicted respectively over time in psychosis, in which the dotted line indicates worse clinical outcome in some patients. This model can be tested in longitudinal designs to clarify the temporal and causal relationship between the different priors.

**Figure 10:**
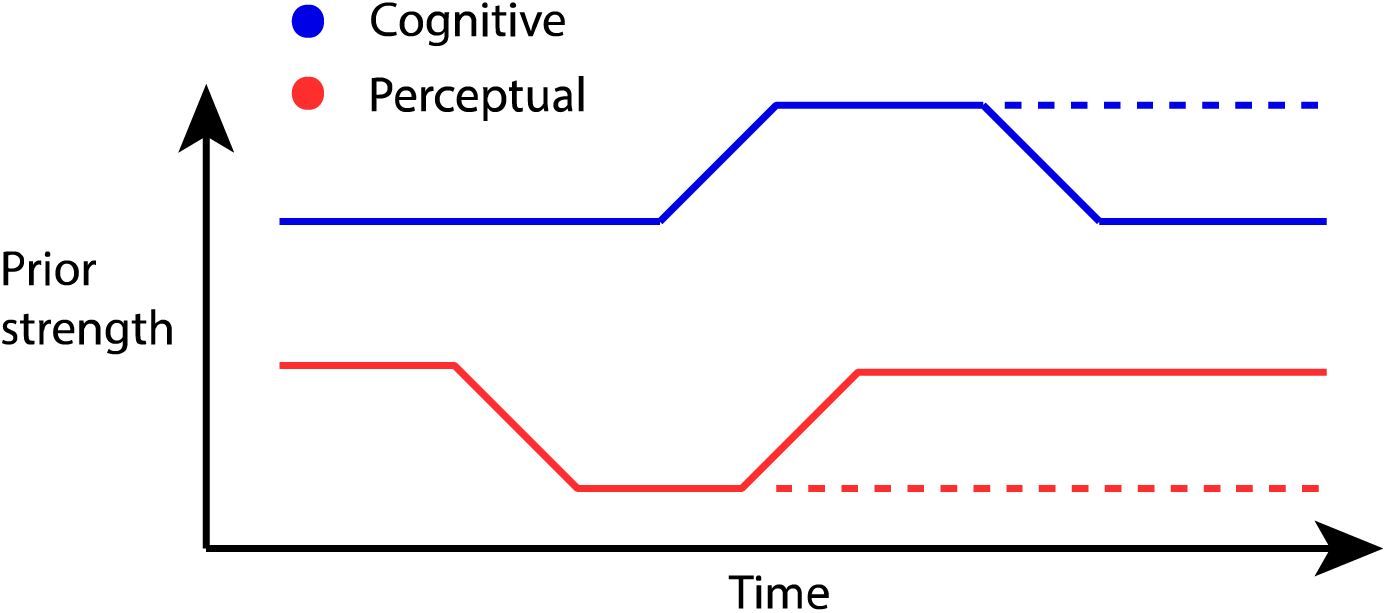
A proposed model for the interaction between different levels of prior over time in psychosis. The early stages of psychosis might be characterized by a weakening of lower-level perceptual priors as indicated by a fall in the lower red line. This causes a shift in the strength of cognitive priors as an attempt to explain the abnormal perceptual experiences, causing positive symptoms of psychosis. This will counter the weakening of lower-level priors A failure to attenuate the weakening of lower-level priors may result in more severe, sustained symptoms as indicated by the dashed lines.

Two previous studies have looked at the McGurk effect in schizophrenia. White et al (2009) found that patients were, on average, less vulnerable to the illusion than controls, with a strong relationship with duration of illness, such that individuals who have been ill for longer were less likely to report a McGurk effect (White et al., 2014). Pearl et al (2009) used a more complex recruitment design and had more mixed results that interacted in a complex fashion with age; the interpretation of their patient results are made challenging given that results in controls interacted with age in an unexpected manner. In these previous studies participants were required to report binary choices on whether they perceived the McGurk effect, whereas we used a staircase procedure to examine the degree of influence that lip-movements have on auditory perception. We did not find a diminishment in the degree that lip-movements influenced auditory perception in psychosis patients. This might relate to differences in methodology, or perhaps to the age difference between our study (mean age 24.9 years) and White’s study (mean age 39.0 years), given that the absence of illusory effect was more marked in White et al’s older patients with longer disease duration. Further studies looked at the ability for schizophrenia patients to use lip-movements to understand written speech, which found aberrancies in schizophrenia, while general lip-reading capabilities remained intact (Myslobodsky et al., 1992; de Gelder et al., 2002; Ross et al., 2007; Pearl et al., 2009; Szycik et al., 2013). Again, the patient groups in these studies consisted of schizophrenia patients who were older and in a more chronic phase than in the present sample, potentially explaining the discrepancy with the present study.

In the present study we have described our effects in terms of an increase or decrease in the influence of prior expectations. However, it should be noted that the present paradigm is not able to directly discern whether a stronger influence of prior expectations in auditory perception is due to a change in the strength of prior or a weakening in the strength of the sensory input. Indeed previous studies have shown impairments in the ability to do simple auditory discrimination tasks in schizophrenia (Javitt et al., 2015). Future studies could utilise simple auditory discrimination tasks to explore whether these effects are driven by these deficiencies or whether they can be separated.

It has been proposed that glutamatergic abnormalities may be prominent in the early stages of psychotic illness (Merritt et al 2016, Kumar et al 2018), and that these may be key in driving pathophysiology of illness, predictive processing dysfunction, and psychopathology (Corlett et al., 2009 & 2011; Sterzer et al 2018). We did not find a significant relationship between glutamate levels in the anterior cingulate cortex and the strength of the perceptual and cognitive priors across all participants. However, in an exploratory analyses, we analysed the groups separately, and here we did find that in the healthy group there was a significant positive relationship between anterior cingulate glutamate levels and cognitive priors, and in the FEP group a significant relationship between glutamate levels and perceptual priors. This relationship between anterior cingulate glutamate levels and perceptual priors in the FEP group is interesting as the correlations suggest that a (sustained) weakening of perceptual priors is particularly relevant to FEP symptomology, and thus glutamate might play a role in having sustained weakened perceptual priors. The absence of a correlation with the cognitive priors might be due to a lack of power, as a successful glutamate scan was only acquired from 14 individuals who had first episode psychosis. We report MRS results uncorrected for multiple comparisons, which should currently be viewed as preliminary. Larger sample size studies on glutamate levels, the strength of perceptual priors in psychosis, and their inter-relation, will be required to confirm (or refute) our results, which should currently be interpreted with caution. A further limitation of our MRS work is the use of a single region, located in the anterior cingulate cortex, from which our glutamate measure is drawn. We do not mean to imply that this is the only region influencing the role of priors in decisions, but until MRS technology matures to allow simultaneous acquisition of neurochemistry measures across the whole brain, a priori region of interest selection will remain the norm.

As with all studies that use the at-risk-mental-state construct, there is an inherent limitation in terms of the inability to prospectively determine whether an at-risk individual will develop a first episode of psychosis. Therefore, future studies would benefit from following up individuals determined to be in the at-risk group, and so explore the predictive validity of a change in the usage of priors. Indeed, longitudinal studies will be required for definitive conclusions about how use of priors relates to illness stage.

Extending this research beyond the field of psychosis, we note that autism has been suggested to also be associated with a weakening of priors, but which usually does not develop into psychotic symptoms (Pellicano et al., 2012; van Boxtel et al., 2013; Lawson et al., 2014), although there are increased rates of psychotic symptoms in autism and other neurodevelopmental disorders (Hussain and Murray 2015; Larson et al., 2017). The difference between schizophrenia spectrum psychosis and autism may lie in the fact that autism presents itself in early childhood, whereas schizophrenia spectrum illness typically develops later in adolescence. The consequence of this is that during the emergence of schizophrenia spectrum psychosis the brain has to explain a changing world, whereas the sensory driven world autism is characterized by presents itself at birth, requiring no changes in the model of the world to form (i.e. no formation of delusional beliefs), yet the experience of being overwhelmed by sensory experiences remains. Future experiments would need to use a longitudinal approach to support this hypothesis, namely that psychosis is preceded by a decrease in the influence of perceptual priors on perception, followed by a normalization accompanied by an increase in higher-level cognitive priors, whereas autism has weakened priors from birth. In order to test such hypotheses, longitudinal paradigms are preferred which require potentially large groups of people. In order to acquire such amounts of data, the possibility of online testing could be considered, for which the present experiments are well adapted too, due to the simplicity of the paradigm and the brief duration of the experiments (10 minutes each).

In conclusion, we found that the influence of perceptual priors might be weakened in the early stages of psychosis but not in the later stages, whereas cognitive priors are strengthened in the later stages but not early stages. We therefore suggest that previous reported inconsistencies in the literature regarding the influence of prior expectations on sensory processing might be due to differences in the origin of the prior expectation and the disease stage. Furthermore, changes in perceptual and cognitive priors might interact with each other throughout the development of psychosis and glutamate might play a mediating role in the process.

## Author roles

JH: conceptualization (lead role), methodology (lead role), software, formal analysis, investigation, writing (original draft preparation, review and editing). FK: formal analysis, supervision, writing (original draft preparation, review and editing). JG: conceptualization, methodology, investigation, formal analysis, writing (review and editing). HT: investigation, formal analysis, writing (review and editing). MM: investigation, methods, formal analysis, writing (review and editing). IG: supervision, project administration, funding acquisition. PCF: conceptualization, project administration, writing (review and editing). GKM, conceptualization, project administration, methodology, supervision (lead role), writing (original draft preparation, review and editing)

## Acknowledgements

We would like to thank Owen Parsons with his help in designing the paradigm, Rachel Anderson and Eleanor van Sprang for their help in data collection, CAMEO staff for help with recruitment, and the participants.

